# Single-cell transcriptomic atlas reveals that vascular tissues orchestrate cell fate transitions to initiate adventitious root formation in tomato

**DOI:** 10.64898/2025.12.27.696651

**Authors:** Shipeng Yang, Cheng Si, Xiaoting Jiang, Xuemei Sun, Cao Shanni, Qiwen Zhong, Ruolin Yang

## Abstract

- Adventitious root (AR) formation enables vegetative propagation critical for agriculture, conservation, and breeding, yet the cellular basis of AR competence in crop species remains poorly characterized. Most mechanistic studies rely on *Arabidopsis*, which exhibits constrained AR capacity compared to species dependent on clonal propagation.
- We constructed a single-cell transcriptomic atlas of tomato (*Solanum lycopersicum*) AR development comprising 7,920 cells across 12 cell types. Regulatory networks were inferred using hdWGCNA and MINI-EX, validated by RNA in situ hybridization and qPCR. Cross-dataset integration with shoot-borne root initiation data enabled developmental staging, while SATURN-based integration of 118,730 vascular cells from eight species permitted evolutionary comparison.
- CytoTRACE analysis revealed that vascular tissues—root stele and phloem—possess the highest developmental potential, supporting a model of latent stem cell-like identity rather than dedifferentiation-based reprogramming. Network analysis identified *LEA3* as a central hub gene constitutively expressed throughout AR development, regulated by the DOF transcription factor *DOF11*. Cross-species integration demonstrated that tomato AR-initiating cells share transcriptomic similarity with woody dicots (*Liriodendron*, *r* = 0.46; *Eucalyptus*, *r* = 0.42) but not *Arabidopsis* (*r* = −0.02), with *LEA3*-containing macrogene weights highest in basal angiosperms and woody species.
- Our findings establish the *DOF11-LEA3* axis as a conserved vascular identity module and suggest that AR competence represents an ancestral program retained in species with active cambial dynamics. These results reframe *Arabidopsis* as a derived rather than representative model for AR biology and identify molecular targets for enhancing vegetative propagation.

## Introduction

The capacity to regenerate organs from differentiated tissues represents one of the most remarkable manifestations of plant developmental plasticity. Unlike animals, where regenerative potential is largely restricted to specific stem cell populations, plants can initiate de novo organogenesis from a wide range of mature cell types—a property that underlies both natural adaptation to environmental damage and agricultural practices such as vegetative propagation (Umeda *et al*., 2021). Adventitious root (AR) formation exemplifies this plasticity: functional roots emerge from stems, leaves, or hypocotyls without traversing embryonic developmental programs, enabling clonal propagation of economically important crops and survival following root system damage (Singh *et al*., 2022; Adem *et al*., 2024). Beyond agriculture, AR competence carries broader significance: it enables ex situ conservation of endangered species through vegetative propagation when seed-based reproduction fails, permits rapid fixation of superior breeding traits without generational segregation, and fundamentally constrains the efficiency of tissue culture and micropropagation systems (Rather *et al*., 2022; Murthy *et al*., 2024).

Despite extensive physiological characterization, the cellular and molecular basis of AR competence remains incompletely understood. The prevailing model posits that AR formation requires dedifferentiation of mature cells followed by re-establishment of a root stem cell niche—a process conceptualized as cellular reprogramming driven by wound signals and auxin accumulation (Xu, 2018). Yet this framework raises fundamental questions: Do AR-competent cells truly undergo complete dedifferentiation, or do they retain latent developmental programs that are activated rather than reconstructed? What distinguishes the subset of cells capable of organogenic fate transitions from their non-responsive neighbors? And critically, are the regulatory mechanisms governing AR initiation conserved across species, or do they reflect lineage-specific adaptations?

Single-cell transcriptomics has begun to illuminate the heterogeneity obscured by bulk tissue analyses. Recent studies have identified transitional cell states during de novo root regeneration, revealing that phloem-associated cells serve as the primary source of new root meristems (Liu *et al*., 2022; Damodaran & Strader, 2024). Time-resolved single-cell profiling has further demonstrated that wound-induced regeneration proceeds through distinct transcriptional phases, with jasmonate, ethylene, and reactive oxygen species pathways activated within hours of tissue excision (Morinaka, 2025). These advances notwithstanding, most single-cell investigations of AR formation have focused on the herbaceous model Arabidopsis thaliana—a species with notably constrained AR capacity compared to woody plants and crops dependent on vegetative propagation.

The reliance on *Arabidopsis* as the primary model may have introduced systematic biases into our understanding of AR biology. Transcription factors functioning as positive regulators of AR formation in woody species paradoxically act as negative regulators in *Arabidopsis* (Liu *et al*., 2025), and the minimal secondary growth in Arabidopsis precludes analysis of cambial contributions to organogenic competence. These observations raise the possibility that Arabidopsis represents a derived rather than ancestral state for AR regulatory networks—a hypothesis with significant implications for translating mechanistic insights to agriculturally relevant species.

Tomato (*Solanum lycopersicum*) offers a compelling alternative system. Unlike *Arabidopsis*, tomato exhibits robust AR formation from stem cuttings, possesses substantial secondary growth capacity, and supports the clonal propagation practices essential to modern horticulture(Zhang, L *et al*., 2025). The recent development of cross-species single-cell integration methods—particularly approaches leveraging protein language models to overcome orthology limitations (Passalacqua & Gillis, 2024; Rosen *et al*., 2024)—now enables systematic comparison of vascular developmental programs across phylogenetically diverse species, potentially revealing whether AR competence reflects conserved ancestral mechanisms or convergent evolution.

Here, we present a comprehensive single-cell transcriptomic atlas of tomato AR development, integrating spatial validation, gene regulatory network inference, cross-dataset developmental staging, and cross-species evolutionary analysis. Our investigation addresses three fundamental questions: (1) What transcriptional features distinguish AR-competent vascular cells, and do they support a model of latent stem cell-like identity versus dedifferentiation-based reprogramming? (2) What regulatory modules coordinate vascular identity with stress protection during the transition from stem tissue to root primordium? (3) Does tomato AR development deploy transcriptional programs conserved with woody species, or does it more closely resemble the constrained regenerative capacity of Arabidopsis? The answers to these questions reframe our understanding of post-embryonic organogenesis and identify molecular targets for enhancing vegetative propagation in crop species.

## Materials and Methods

### Preparation of AR Samples for scRNA-Seq

Heinz 1706 tomato plants (approximately 1.2 m in height) were selected for AR induction. Lateral stems (∼10 cm) were excised at the axillary node and, within 1 h of detachment, transferred to a hydroponic cultivation system in a controlled growth chamber. The culture water (resistivity: 18 MΩ·cm) was provided by a Millipore unit, and illumination was supplied by Philips GreenPower LEDs at 800 lx for 12 h, followed by 12 h of darkness. Each excised stem was secured on a floating foam board such that the bottom 1 cm was submerged. After 55 h, AR primordia emerged at the stem base; roots ranging from ∼0.1 to 1 cm in length were harvested in bulk, flash-frozen in liquid nitrogen, and stored at −70 °C until further use.

### Single-Cell Library Construction and Sequencing

Protoplasts were isolated from AR tissue using an optimized enzymatic digestion protocol (Methods S1). A small aliquot of the cell suspension was stained with 0.4% Trypan Blue and counted on a Countess® II Automated Cell Counter to ensure a viable cell concentration of ∼1,000–2,000 cells/μL. Each sample was then encapsulated with 10x Genomics gel beads and reverse**-**transcription reagents, forming oil-partitioned GEMs (Gel Beads**-**in**-**Emulsions). Within each GEM, gel beads were lysed to release barcode sequences, enabling cDNA synthesis and indexing. Subsequently, the emulsions were broken, and the pooled cDNA was PCR**-**amplified. Following fragmentation to ∼200–300 bp, standard Illumina adaptors (P5, P7, and sample indices) were added, and the resulting libraries were amplified via PCR. Final libraries were sequenced on an Illumina NovaSeq 6000 using PE150 protocol by Gene Denovo Biotechnology (Guangzhou, China).

### Data Preprocessing and Quantification

Raw sequencing reads underwent initial quality control (QC) and demultiplexing with the Cell Ranger software suite (10x Genomics). Low**-**quality reads were discarded, and remaining reads were aligned to the tomato reference genome SL4.0 (https://solgenomics.net/organism/Solanum_lycopersicum/genome) using the STAR aligner (v2.7.10b) (https://github.com/alexdobin/STAR). Cell Ranger (version 2.0, 10x Genomics) annotated reads to specific genes, corrected unique molecular identifiers (UMIs), and generated an unfiltered feature**-**barcode matrix. Putative cells were distinguished from non**-**cell barcodes according to Cell Ranger’s default algorithm, providing a preliminary cell count for downstream analysis.

### Seurat-Based Analysis

All downstream analyses were performed in R (v4.1.2) using the Seurat package (v4.0.5). Single-cell data were loaded with minimum cutoffs of three cells per gene and 200 genes per cell. Quality control filtering retained cells expressing 200–4,000 genes and <6% mitochondrial content. Data were log-normalized with a scale factor of 10,000, and the top 2,000 most variable genes were selected for downstream analysis. All genes were scaled prior to dimensionality reduction by principal component analysis (PCA). The first 30 principal components were used to construct a shared nearest neighbor (SNN) graph (k = 90), and cells were clustered at resolution 0.8. The resulting 12 distinct subpopulations were annotated according to known marker genes and differential expression profiles, then visualized using Uniform Manifold Approximation and Projection (UMAP).

### Marker Gene Identification and In Situ Hybridization

To define the cell types of *Solanum lycopersicum* ARs, we first leveraged reported markers from *Arabidopsis thaliana* (Denyer *et al*., 2019; Zhang *et al*., 2019; Dorrity *et al*., 2021), *Oryza sativa* (Zhang *et al*., 2021), and *Triticum aestivum* (Zhang *et al*., 2023), mapping their homologs in *S. lycopersicum* via orthology. We then employed the SPMarker pipeline (Yan *et al*., 2022), which utilizes machine learning–based feature importance (e.g., SVM, random forest, Shapley additive explanations) to identify discriminative cell-type markers. This two-pronged strategy—marker homology plus SPMarker—facilitated robust annotation of putative cell clusters and guided differential expression analyses. Gene-specific probes were synthesized following the Dig Northern Starter Kit (Roche) protocol. Young ARs were harvested at a similar developmental stage, fixed in 50% formaldehyde–acetic acid–ethanol (FAA) at 4 °C for 24 h, then paraffin-embedded and sectioned to 10 µm. Deparaffinized sections underwent sequential xylene/ethanol washes and were boiled in antigen-retrieval solution. After proteinase K digestion (20 µg mL**⁻**¹, 37 °C, 10 min), slides were prehybridized at 37 °C (1 h) and hybridized overnight at 42 °C with digoxigenin-labeled RNA probes. Post-hybridization washes were followed by microscopic visualization (Nikon Eclipse 80i), confirming the spatial expression patterns of these newly identified AR-specific markers.

### Functional Annotation with CellFunTopic

To gain insights into cluster-specific biological processes, we used the CellFunTopic package in R (v4.1.2). After the standard Seurat-based filtering, normalization, and PCA, the top 10 principal components were selected, and unsupervised clustering at resolution 1.0 identified distinct cell groups. Gene set enrichment analysis (GSEA) was performed with Gene Ontology (GO) terms as the primary database to calculate enrichment scores and FDR-adjusted p-values for each cluster. Significant pathways were visualized through pathway heatmaps, hierarchical clustering of pathway relationships, and UMAP embeddings annotated with pathway enrichment scores. Cluster-level correlations based on Pearson’s or Jaccard coefficients of enriched GO terms were computed to reveal functional similarities among cell populations. Distribution patterns of selected pathways were displayed in 2D embeddings, enabling comprehensive functional annotation of each cluster.

### Pseudotime and Differentiation Trajectory Analysis

To explore the hierarchical progression of tissue-specific lineages during AR formation, we first used CytoTRACE (v0.3.3) to quantify each cell’s relative differentiation potential (Gulati *et al*., 2020). A gene-count matrix was extracted from the Seurat object and filtered to retain transcripts with nonzero counts in at least five cells. CytoTRACE generated a continuous differentiation score for each cell, which was visualized alongside cluster annotations and dimensionality-reduction embeddings. For trajectory inference, we employed two complementary approaches. First, Monocle3 was used to order cells along a pseudotime axis, delineating transitions from vascular-associated precursors to more specialized cortical and epidermal derivatives. Second, lineage reconstruction was performed with Slingshot (v2.4.0), connecting putative progenitor-like populations with terminal cell fates to reveal branching lineages with corresponding pseudotime values (Street *et al*., 2018). We further identified differentially expressed genes along each lineage using tradeSeq (v1.10.0), fitting negative binomial generalized additive models to capture dynamic gene expression patterns (Van den Berge *et al*., 2020). Significant gene sets exhibiting pseudotime-dependent expression changes were identified, delineating distinct modules of vascular, ground, and epidermal cell differentiation.

### Co-expression Network Construction and Analysis with hdWGCNA

We employed the hdWGCNA pipeline (v0.3.00) to identify and characterize co-expression modules from our scRNA-seq dataset (Morabito *et al*., 2023). First, highly variable genes were selected and relevant parameters were stored for network construction. To mitigate technical noise and stabilize expression signals, cells were grouped into metacells (k = 25), followed by normalization of aggregated counts. An optimal soft-power threshold was estimated to approximate scale-free topology. The topological overlap matrix (TOM) was computed to define co-expression modules in a signed network. Each module’s eigengene was derived, and intramodular connectivity was quantified to identify hub genes. Module assignments were refined and examined through dendrogram visualization and network topology plots. For functional characterization, enrichment analysis of module gene sets was performed using Gene Ontology annotations, thereby associating each module with overrepresented biological processes. Phylogenetic trees of hub gene *LEA3* were constructed using SHOOT (Emms & Kelly, 2022).

### MINI-EX Analysis of Single-Cell Gene Regulatory Networks

We used MINI-EX (Motif-Informed Network Inference based on single-cell EXpression data) to infer and prioritize cell-type-specific gene regulatory networks (GRNs) in tomato ARs (Ferrari *et al*., 2022). The pipeline was provided with a curated TF annotation file and the single-cell gene-by-cell expression matrix in raw counts. The analysis workflow included: (1) construction of an initial expression-based regulatory network via GRNBoost2, (2) filtering of inferred TF–target gene interactions by TF binding-site (TFBS) enrichment using both TF-specific and TF-family motif information, and (3) assignment of validated regulons to specific cell clusters through enrichment of cluster upregulated genes. TFs sufficiently expressed in corresponding cell clusters were retained as bona fide regulators. To rank candidate regulators, MINI-EX applied network centrality metrics (betweenness, closeness, degree) in conjunction with cluster-level differential expression and functional enrichment. Weighted Borda scores integrating expression specificity, centrality measures, and functional annotations were computed for each regulon. Phylogenetic relationships of key regulator *DOF11* were inferred using SHOOT (Emms & Kelly, 2022).

### Promoter Cis-Regulatory Element Analysis

Promoter sequences (2,000 bp upstream of transcription start sites) were extracted from the tomato SL4.0 genome (ITAG4.0 annotation). Auxin-responsive elements (AuxRE: TGA-element, AuxRR-core, AuxRE-core) and ABA-responsive elements (ABRE and variants) were identified using exact sequence matching on both strands based on PlantCARE database motifs (Lescot *et al*., 2002). ABRE:AuxRE ratios were calculated to assess the relative dominance of ABA-versus auxin-responsive regulation (Methods S2).

### qPCR Validation of Hub Gene Expression

To validate hormone responsiveness of hub genes, tomato seedlings were treated with IAA (100 **μ**M), ABA (100 **μ**M), or their combination, with samples collected at seven time points (0–144 h). Total RNA was extracted using TRIzol, reverse-transcribed using PrimeScript RT Kit (Takara), and quantified using TB Green-based qPCR on a LightCycler 480 II system. Relative expression was calculated using the 2^−**ΔΔ**Ct^ method with *SlActin* as reference. Statistical significance was determined by one-way ANOVA with Tukey’s HSD post-hoc test (P < 0.05).

### Primer sequences are provided in Supplementary Table S8 (Methods S3). Cross-Dataset Integration and Developmental Trajectory Analysis

The SlAR dataset (7,920 cells) was integrated with a published shoot-borne root (SBR) dataset (Omary et al., 2022; GSE159055; 2,671 cells) across 20,410 shared genes. Seven batch correction methods were benchmarked using scib-metrics (Luecken *et al*., 2022), with ComBat (Johnson *et al*., 2007) selected based on highest total score (0.678), balancing batch mixing (0.571) and biological conservation (0.750). A composite developmental score was computed using six early-stage markers and 14 late-stage markers, positioning cells along the initiation-to-maturation continuum (Methods S4).

### Bulk RNA-Seq Analysis

Adventitious root samples were collected at four developmental time points (2, 12, 48, 96 h; three replicates each) and subjected to strand-specific RNA-seq using NEBNext Ultra II kit on Illumina NovaSeq 6000 (PE 2×150 bp). Reads were aligned to tomato SL4.0 using HISAT2 v2.2.1 and quantified using featureCounts v2.0.3. Expression of 28 key genes (hub genes, MINI-EX TFs, SBR regulators) was analyzed using CPM normalization and Z-score transformation (Methods S5).

### SATURN Cross-Species Single-Cell Integration

Single-cell transcriptomes from eight plant species representing diverse phylogenetic lineages were integrated using SATURN (Rosen *et al*., 2024), which employs ESM-2 protein language model embeddings (Lin *et al*., 2023) to enable cross-species gene expression comparison. Published datasets were obtained from public repositories: *Arabidopsis thaliana* stem (GSE226097; 48,479 cells) (Lee *et al*., 2025), *Eucalyptus grandis* stem xylem (GSE180121; 7,889 cells) (Chen *et al*., 2024), *Glycine max* stem (GSE270392; 8,671 cells) (Zhang, X *et al*., 2025), *Liriodendron chinense* stem (GSE180121; 2,977 cells) (Chen *et al*., 2024), *Oryza sativa* stem (GSE232863; 19,740 cells) (Wang *et al*., 2025), *Populus trichocarpa* stem (PRJNA703312; 25,638 cells) (Li *et al*., 2021), and *Trochodendron aralioides* stem (GSE180121; 3,131 cells) (Chen *et al*., 2024). From tomato, only vascular-origin cells (Root stele, Protoxylem, Phloem; 2,205 cells) were retained to ensure tissue comparability with stem vascular tissues from other species. SATURN constructed 500 macrogenes enabling cross-species expression comparison. Cells were clustered into eight pan-cell types, and cross-species expression similarity was computed using Pearson correlation. The LEA3-containing macrogene (MG_420) was identified for comparative analysis (Methods S6).

### OrthoFinder Ortholog Identification

Protein sequences from eight species were processed using OrthoFinder v2.5.4 to identify orthogroups (Emms & Kelly, 2019). The analysis employed DIAMOND for all-vs-all BLASTP searches with default e-value threshold (1e-3), MAFFT for multiple sequence alignment, and IQ-TREE for gene tree inference. Species tree was inferred using the STAG algorithm. For *LEA3*, orthogroup OG0000911 containing 32 genes across all eight species was identified. For *DOF11*, orthogroup OG0000302 containing 45 genes was recovered. Overlap between OrthoFinder orthogroups and SATURN macrogenes was assessed to validate cross-method consistency: *LEA3* showed 31.3% overlap (10/32 genes), while *DOF11* showed 8.9% overlap (4/45 genes), reflecting the complementary nature of sequence-based orthology and embedding-based functional similarity approaches.

## Results

### Construction of Single-Cell Transcriptome Atlas of the Developing Tomato AR

We generated a single-cell transcriptome atlas of tomato AR development following stem excision by combining systematic sampling with an optimized protoplast isolation workflow that preserves cell viability and transcriptional integrity **(Methods S1; Fig. 1a)**. Using the 10x Genomics Chromium platform, we profiled 10,363 cells (393,640,040 reads), capturing a median of 2,207 genes and 4,405 UMIs per cell **(Table S1)**. After alignment and stringent quality control (**Table S2**), 7,920 high-quality cells expressing 25,804 genes were retained for downstream analyses, with vascular-associated populations representing the largest fractions (e.g., root stele: 1,144 cells; **Table S3**).

**Figure 1.**
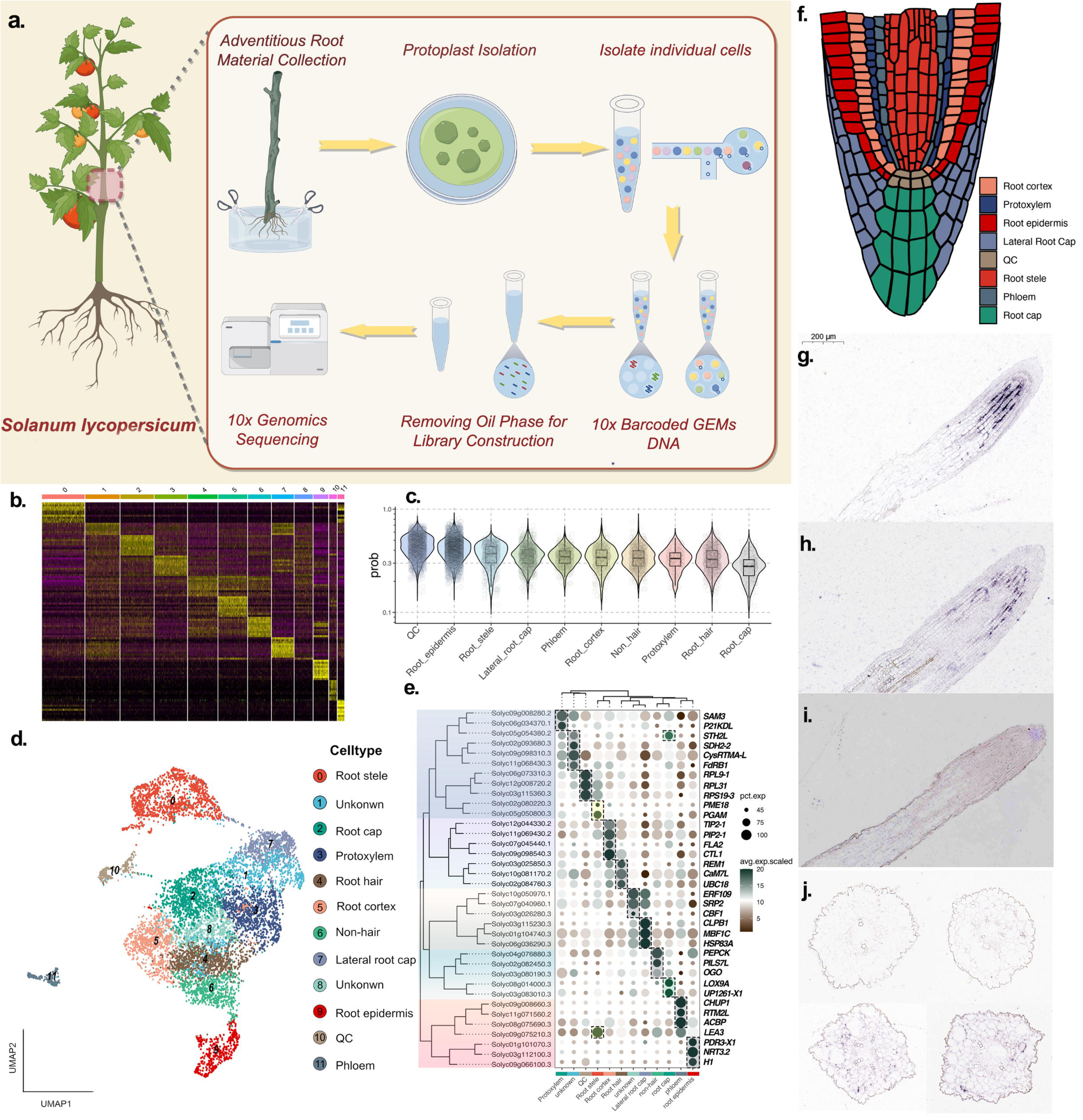
Construction and validation of a single-cell transcriptome atlas of developing tomato ARs. **(a)** Workflow for tomato AR single-cell RNA sequencing, including protoplast isolation and 10x Genomics library preparation. **(b)** Heatmap of cluster-specific gene expression across identified cellular populations. **(c)** Violin plots of SPMarker prediction scores for major AR cell types. **(d)** UMAP visualization of 12 clusters colored by inferred cell type identity. **(e)** Dot plot showing expression of representative marker genes across clusters. **(f)** Schematic illustration of AR tissue organization. (g-j) RNA in situ hybridization showing spatial expression of *LEA3* (Solyc09g075210.3; g,j), *CHUP1* (Solyc09g008660.3; h), and *UP1261-X1* (Solyc03g083010.3; i) in transverse sections along the proximal-distal axis of developing ARs. Scale bar, 200 μm.

We next optimized clustering to resolve discrete transcriptional states that could be mapped to canonical root tissues. A Seurat resolution of 0.8 yielded robust separation while preserving biological structure (**Fig. S1**), defining 12 transcriptionally distinct cell populations (**Fig. 1b,d**). Cell identities were assigned by integrating marker-based annotation with SPMarker supervised predictions, which showed high confidence for the quiescent center (QC), epidermis, and stele (**Fig. 1c; Fig. S2**).

Notably, root cap–associated clusters exhibited comparatively lower classifier confidence, motivating orthogonal spatial validation by RNA in situ hybridization.

The resulting UMAP embedding separated major root tissue populations—including root stele, protoxylem, cortex, epidermis, root cap, lateral root cap, and phloem—each characterized by distinct transcriptional signatures (**Fig. 1d**). Differential expression analysis identified cell-type-specific markers that robustly distinguished these populations (**Fig. 1e; Table S4**), and their spatial organization was consistent with canonical root architecture (**Fig. 1f**). RNA in situ hybridization confirmed the predicted localization of representative markers across tissues with different prediction confidence (**Fig. 1g–j**). *LEA3* (Solyc09g075210.3) displayed a proximal–distal gradient within the stele, with higher signal near the root tip and reduced signal proximally (**Fig. 1g,j**). The phloem marker *CHUP1* (Solyc09g008660.3) localized to peripheral stele domains (**Fig. 1h**), whereas the root cap marker *UP1261-X1* (Solyc03g083010.3) was restricted to the root tip (**Fig. 1i**). Collectively, these analyses assigned high-confidence identities to the 12 clusters and experimentally confirmed the predicted spatial localization of representative stele, phloem, and root cap markers (**Fig. 1g-j**).

### Fine-scale functional mapping reveals cell type-resolved pathway organization and coordinated hormone-associated programs

Building on the spatially validated cell-type framework (**Fig. 1**), gene set enrichment analysis (GSEA) was applied to delineate how functional programs are partitioned across tomato AR tissues (**Fig. 2a,b**). QC, lateral root cap, and cortex displayed the largest sets of uniquely enriched pathways, consistent with their specialized roles in niche maintenance and environmental sensing. In QC cells, protein heterodimerization activity and chromatin-related terms were among the most significantly enriched categories (**Fig. 2b**), indicating a regulatory context compatible with integrating diverse signaling inputs. Cortex-specific enrichment of water transport and dehydration response pathways aligned with evidence that hydrotropic sensing is associated with elongating cortex tissues. In contrast, auxin-activated signaling was preferentially enriched in vascular-associated tissues (stele and phloem) (**Fig. 2b**), consistent with reported auxin accumulation and transport routes during AR initiation(Guan *et al*., 2019; Zhang *et al*., 2021; Hernández-García *et al*., 2024). Enrichment of cell differentiation–related pathways in the root stele further supported chromatin-level reprogramming during this process. The topic–cluster network summarized functional relatedness among tissues through shared pathway structure (**Fig. 2b**).

**Figure 2.**
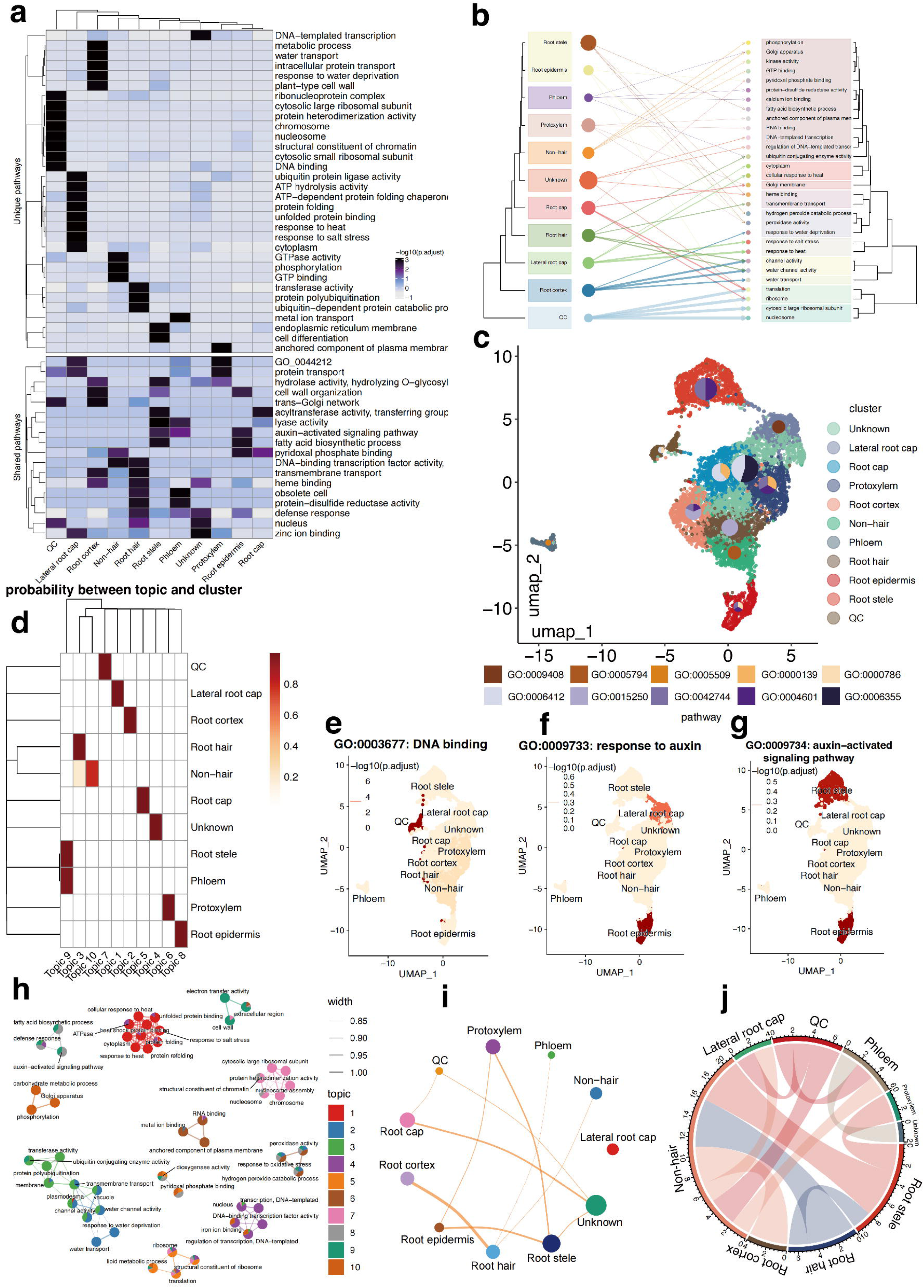
Cell Type-Specific Functional Organization and Pathway Networks in Tomato ARs Development. **(a)** Heatmap showing uniquely enriched (top) and shared (bottom) pathways across cell types; color indicates - log10(adjusted *P* value). **(b)** Topic-cluster network illustrating functional relationships among cell types based on enriched pathways; edge width indicates pathway sharing. **(c)** GSEA-UMAP visualization of cell type distributions with embedded GO terms for the top enriched pathways. **(d)** Topic-cluster probability matrix summarizing cell type-specific functional specialization. **(e)** UMAP projection of DNA binding (GO:0003677) enrichment. **(f)** UMAP projection of response to auxin (GO:0009733) enrichment. **(g)** UMAP projection of auxin-activated signaling (GO:0009734) enrichment. **(h)** Functional module network of pathway interactions across topics. **(i)** Cell-type relationship network based on functional annotations. (j) Circos plot summarizing global topic-function enrichment and cluster associations.

To resolve spatial functional domains within the atlas, GSEA-UMAP analysis of the top enriched pathways highlighted clear tissue-associated patterns (**Fig. 2c**). Peroxidase activity (GO:0004601) was highly enriched in root epidermis, root cortex, and root hair cells, consistent with a redox-associated program implicated in modulating IAA homeostasis through hydrogen peroxide-mediated oxidation. Topics defined by shared biological processes generated a probability matrix capturing the functional organization of root tissues (**Fig. 2d**). Vascular tissues (root stele and phloem) showed high probability within Topic 9, whereas root hair and non-hair populations exhibited distinct profiles in Topics 3 and 10, respectively (**Fig. S5**).

Representative pathways further revealed tissue-restricted enrichment landscapes. DNA binding (GO:0003677) was strongly enriched in QC cells (**Fig. 2e**). Response to auxin (GO:0009733) was enriched in lateral root cap and root epidermis (**Fig. 2f**), whereas auxin-activated signaling (GO:0009734) showed a more specific enrichment pattern in root stele and root epidermis (**Fig. 2g**), consistent with localized auxin redistribution across vascular and peripheral tissues during AR development. This organization of auxin-associated programs parallels reported ROS dynamics during AR induction (Huang *et al*., 2020; Templalexis *et al*., 2022).

Network-level visualization of pathway interactions resolved coordinated functional modules across topics, with closely related pathways clustering into coherent groups (**Fig. 2h; Fig. S6**). Cell-type relationship analysis further revealed structured associations between anatomically adjacent tissues, including root cortex, root hair, and non-hair populations in the outer layers, and root epidermis, root stele, and protoxylem in inner layers (**Fig. 2i**). A circos representation summarized the global landscape of topic-function enrichment and cluster associations (**Fig. 2j**). Collectively, these analyses delineate a cell type-resolved functional organization of tomato AR tissues, with hormone-associated programs exhibiting domain-specific enrichment across vascular and peripheral compartments.

### Pseudotime Analysis Reveals Hierarchical Tissue Differentiation During ARs Formation

To complement the cell type-resolved functional landscape (**Fig. 2**) with a developmental ordering, trajectory inference and potency estimation were used to arrange AR cells along a continuous differentiation axis. Hierarchical clustering of pseudotime-ordered expression profiles resolved three transcriptional modules with distinct temporal peaks and tissue associations (**Fig. 3a**). Module 1, characterized by genes peaking at intermediate pseudotime stages, was enriched for protoxylem, non-hair, and cortex markers (*STH2L*, *SAM3*, *PEPCK*, *PIP2-1*, *TIP2-1*). Module 2, predominantly active at later pseudotime stages, featured lateral root cap and epidermis markers (*MBC1F*, *HSP83A*, *NRT3.2*, *PDR3-X1*), consistent with acquisition of protective and nutrient-uptake functions. By contrast, Module 3 exhibited early pseudotime expression patterns enriched for phloem and stele markers (*ACBP*, *LEA3*), consistent with early establishment of vascular identity and auxin-associated programs during AR initiation.

**Figure 3.**
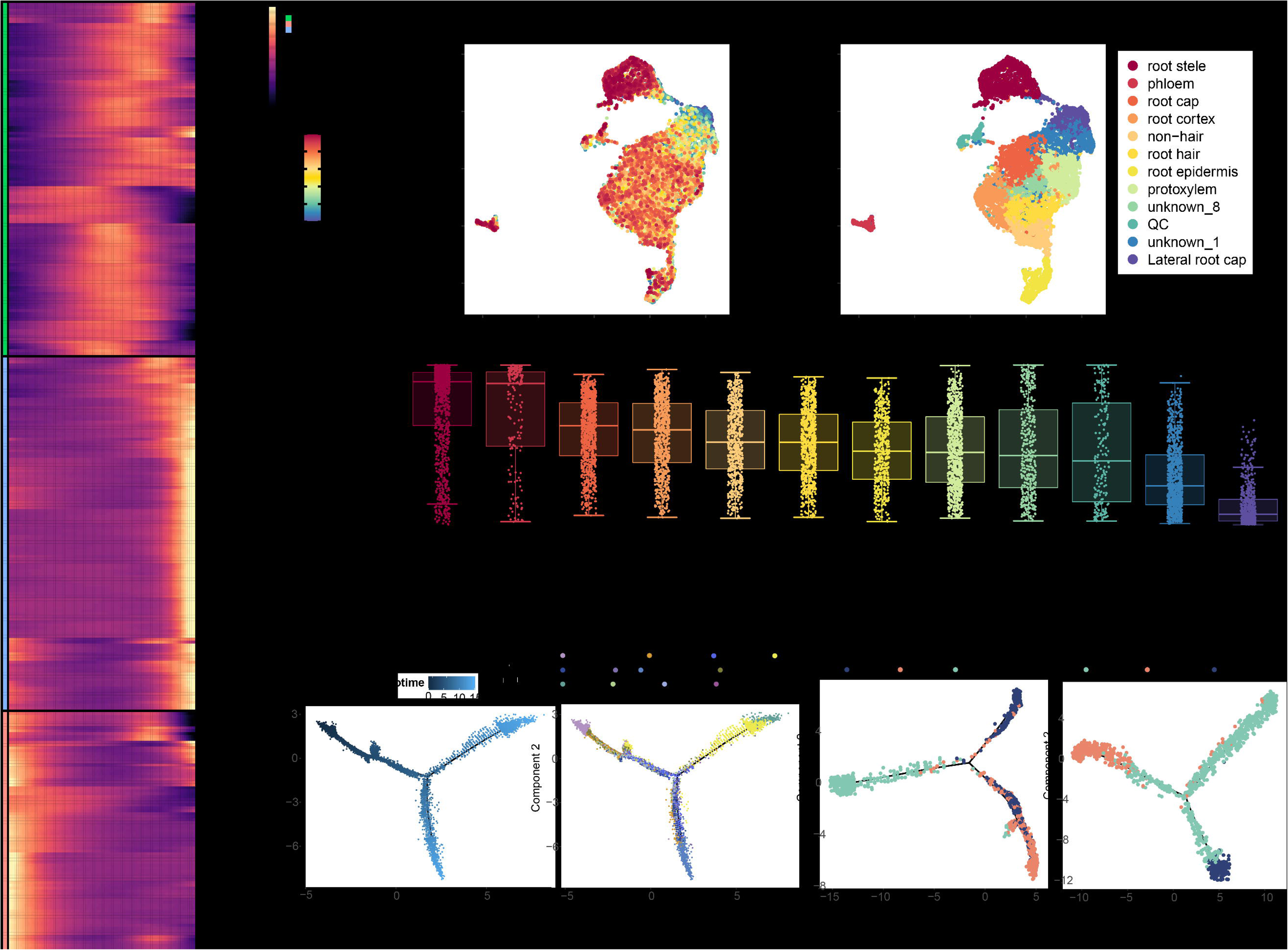
Pseudotime and CytoTRACE Analysis of Tissue Differentiation During Tomato AR Formation. **(a)** Heatmap of pseudotime-ordered expression for module-associated marker genes; rows indicate genes and columns indicate cells ordered along pseudotime. **(b)** UMAP colored by CytoTRACE score, reflecting inferred developmental potential. **(c)** UMAP colored by annotated cell types. **(d)** CytoTRACE ordering scores across cell types, highlighting early-diverging vascular-associated populations (stele and phloem). **(e)** Pseudotime landscape projected onto the embedding. **(f)** Pseudotime trajectories colored by major cell types. **(g)** Epidermal lineage highlighting divergence of root hair from non-hair cells. **(h)** Vascular lineage highlighting protoxylem differentiation along the vascular route.

Pseudotime structure and CytoTRACE scores converged on a hierarchical differentiation framework in which vascular-associated populations diverge early and subsequently give rise to ground-tissue and epidermal derivatives (**Fig. 3b-f**). This topology resembles concentric tissue patterning described in embryonic root development (Denyer *et al*., 2019; Ryu *et al*., 2019; Zhang *et al*., 2021), yet here it is deployed in a post-embryonic context where AR founder cells originate from cambium-or phloem-parenchyma–derived transition zones regulated by AIL/PLT modules (Druege *et al*., 2019; Omary *et al*., 2022; Eswaran *et al*., 2024). Consistently, stele and phloem exhibited the highest CytoTRACE ordering scores (**Fig. 3d**), indicating elevated developmental potential relative to more differentiated populations.

Trajectory geometry further resolved branching relationships between major tissue lineages (**Fig. 3e-h**). Cortex and epidermal precursors mapped to a shared early trajectory before bifurcating toward specialized fates (**Fig. 3f**). Within the epidermal lineage, root hair cells diverged from non-hair epidermal cells (**Fig. 3g**), whereas protoxylem differentiated along the vascular developmental route (**Fig. 3h**). Together, the inferred trajectories define an ordered sequence of tissue differentiation in tomato ARs, with early-emerging vascular-associated states preceding ground and epidermal specialization.

### Co-expression network analysis identifies hormone-responsive hub modules linking AR development with stress-associated programs

The trajectory framework highlighted early-emerging vascular-associated states (**Fig. 3**), prompting an assessment of coordinated gene programs and candidate hubs operating across tissues. High-dimensional weighted gene co-expression network analysis (hdWGCNA) of 1,741 highly variable genes (**soft power = 7; Fig. S7**) resolved 11 co-expression modules with pronounced tissue specificity (**Fig. 4a,b**). Multiple modules were enriched in root stele populations (modules 4, 6, 7, 8, and 11), whereas epidermal-associated signal was prominent in modules 1 and 4 (**Table S5**). Module specificity was supported by uCell scoring (**Fig. S8**), and GO enrichment further distinguished functional programs, including stem cell maintenance–related terms in QC and hormone response–related pathways in root stele (**Fig. 4c**).

**Figure 4.**
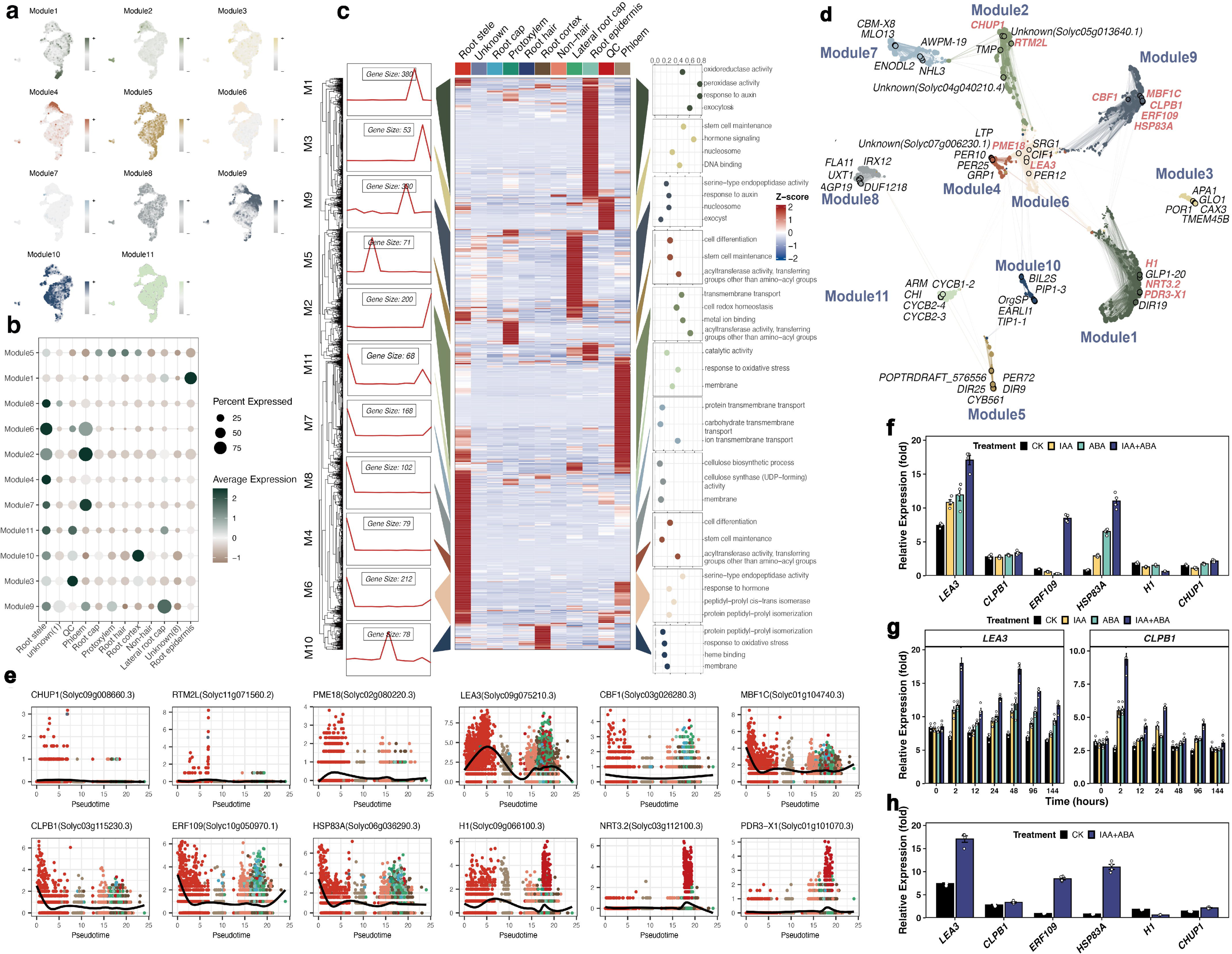
Co-expression network analysis resolves modular programs and nominates hormone-responsive hub regulators during tomato AR development. **(a)** UMAP projections of hdWGCNA module scores showing module-specific activity across AR tissues. **(b)** Dot plot summarizing module prevalence and expression intensity across cell types (dot size, fraction of cells; color, mean module score). **(c)** Heatmap of module dynamics along pseudotime with representative temporal patterns; GO enrichment terms are shown to the right. **(d)** Network topology highlighting hub genes within key modules. **(e)** Pseudotemporal expression profiles of representative stress-associated genes, including *LEA3*, *CLPB1*, and *ERF109*. **(f)** qPCR validation of hub gene expression at 48 h under CK, IAA, ABA, and IAA+ABA treatments. **(g)** Time-course expression dynamics of *LEA3* and *CLPB1* across 0-144 h. **(h)** *LEA3* induction under IAA+ABA relative to control at 48 h. Data are shown as mean ± SEM (n = 4 biological replicates).

Network topology prioritized *LEA3* (module 6; stele-enriched) and *CLPB1* (module 9; lateral root cap–enriched) as central hubs, each peaking at early pseudotime (**Fig. 4d**). Additional stress-associated genes including *ERF109* showed coordinated pseudotemporal expression with these hubs within their respective module contexts (**Fig. 4e**), consistent with a coupled developmental and stress-associated transcriptional program during AR formation.

Promoter cis-element analysis provided a mechanistic framework for differential hormone responsiveness of these hub genes (**Fig. S9; Table S6**). The *LEA3* promoter contained four ABA-responsive elements (ABREs) and one auxin-responsive element (AuxRE) (ABRE:AuxRE = 4.0), whereas the *CLPB1* promoter harbored five ABREs and two AuxREs (ABRE:AuxRE = 2.5), together with a TGA element implicated in stress signaling (**Table S7**). Consistent with these promoter architectures, qPCR profiling of tomato stem explants treated with IAA, ABA, or IAA+ABA across 0-144 h (**Table S8**) showed that, at 48 h-coinciding with AR primordium emergence-LEA3 exhibited the strongest induction among the tested hub genes, with IAA+ABA eliciting significantly higher expression than either single treatment (**Fig. 4f**). Temporal trajectories further distinguished hub dynamics (**Fig. 4g**): *LEA3* maintained an elevated baseline (7.2-fold) and showed additional enhancement under IAA+ABA at 2 h and 48 h, whereas *CLPB1* displayed a rapid early response at 2 h followed by a gradual decline. At 48 h, IAA+ABA co-treatment increased LEA3 expression by ∼2.4-fold relative to control (**Fig. 4h**), aligning sustained *LEA3* expression with an ABA-biased cis-regulatory landscape and contrasting it with a more transient induction pattern for *CLPB1*.

Multiple orthogonal results converged on *LEA3* as a central vascular feature of tomato AR development, supported by its module-central position in the co-expression network and sustained induction under combined ABA and auxin treatments (**Fig. 4**). Phylogenetic reconstruction added an additional layer of context: tomato *LEA3* homologs formed a tight *Solanaceae* sister pair embedded within a broadly diversified eudicot LEA clade (**Fig. S10**), whereas *CLPB1* showed a comparatively conserved placement closer to *Brassicaceae*/*Arabidopsis* homologs (**Fig. S11).**

### Cell Type-Resolved Regulatory Networks Uncover Dynamic Transcriptional Programs in AR Formation

The functional and developmental stratification of the AR atlas (**Figs. 2-3**) implies that transcriptional control should be unevenly distributed across tissues, with regulatory complexity concentrating in founder-associated vascular domains. Consistent with this, motif-informed network inference at single-cell resolution resolved distinct regulatory modules across clusters (**Fig. 5a-d; Fig. S12**).

**Figure 5.**
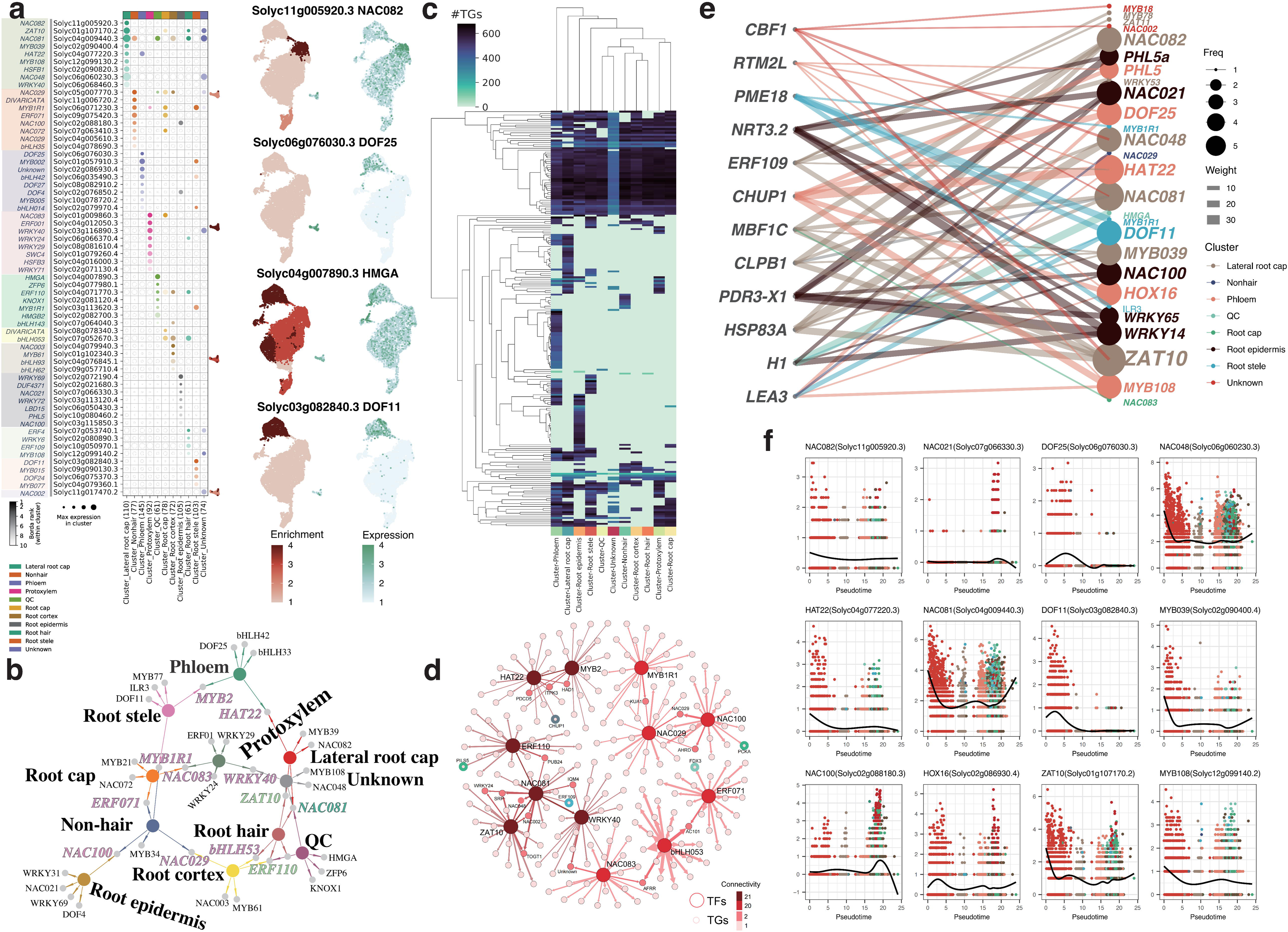
Cell type-resolved regulatory networks identify vascular regulatory centers and nominate DOF11 as a leading upstream regulator of LEA3. **(a)** MINI-EX inference of TF activity across clusters shown as dot plot and UMAP projection for representative regulators. **(b)** Tissue-associated regulatory modules highlighting shared and cluster-specific TF programs. **(c)** Comparison of inferred regulatory complexity across clusters, including target-gene (TG) abundance and hierarchical relationships; vascular-associated clusters show elevated TG counts relative to other populations. **(d)** Network visualization of representative TF-TG interactions; node size reflects regulatory weight and edge thickness reflects interaction strength. **(e)** Bipartite TF-TG network integrating MINI-EX predictions with hdWGCNA hub genes, highlighting DOF-centered regulation of module-6 targets including *LEA3*. **(f)** Pseudotime dynamics of representative TFs, showing early induction of *DOF11*/*DOF25* during AR initiation.

Among these, NAC family factors were broadly enriched, with particularly prominent representation in root cap and lateral root cap populations (**Fig. 5a; Table S10**), indicating extensive TF activity in boundary and protective tissues. In contrast, vascular-associated clusters were distinguished by dominant DOF regulators: *DOF25* and *DOF11* ranked among the highest-weight candidates within the stele-phloem axis (**Table S9**), accompanied by cross-tissue regulators including *ZAT10*, *NAC081*, and *ERF110* (**Fig. 5b; Table S10**). Network architecture further supported this asymmetry: stele-and phloem-associated clusters-linked to AR initiation-exhibited markedly higher numbers of inferred targets than other clusters, whereas unassigned clusters showed minimal regulatory complexity (**Fig. 5c; Table S11**). Dendrogram structure mirrored this pattern, with regulatory differentiation most pronounced within vascular populations (**Fig. 5c**), consistent with these tissues acting as primary regulatory centers during AR formation.

Integration of inferred TF-target relationships with the co-expression framework further refined a focused vascular regulatory axis. DOF-centered interactions were strongly enriched toward the stele-enriched module-6 hub set, with *LEA3* emerging as a top-connected target within this interface (**Fig. 5e; Table S12**). Within the ranked regulator list, *DOF11* showed the strongest overall support as an upstream regulator of *LEA3*, achieving the highest global Borda score and interaction weight in the vascular context (Borda = 570; weight = 46.4), exceeding *DOF25* and all other candidates (**Table S12**). Developmental ordering reinforced this prioritization: *DOF11* and *DOF25* were sharply induced in the earliest pseudotime interval (0-5) (**Fig. 5f**), positioning *DOF11* as an early-acting driver converging on the *LEA3*-associated program.

### Cross-dataset integration quantifies developmental divergence during post-embryonic adventitious root formation

The LEA3-centered vascular program identified in the SlAR atlas raised a staging question: do these maturation-associated signatures distinguish SlAR from transcriptomes capturing earlier post-embryonic root initiation? To address this, SlAR (7,920 cells) was integrated with a published tomato shoot-borne root (SBR) initiation atlas derived from stem pericycle (2,671 cells; Omary et al., 2022; GSE159055) (**Table S13**). Although both datasets represent post-embryonic root organogenesis, they were generated from distinct developmental windows-SBR emphasizing early primordium specification and SlAR capturing more differentiated tissue states.

After filtering to 20,410 shared genes, integration performance was benchmarked across seven batch-correction strategies using scib-metrics (**Fig. 6a; Table S14**). ComBat achieved the strongest overall balance between batch mixing and biological signal preservation (total score = 0.65) and was used for downstream joint embedding (**Fig. 6b, c**). In the integrated space, SlAR resolved 12 mature cell types dominated by root stele (28.0%), root cap (16.7%), and protoxylem (15.0%) (Table S15), whereas SBR comprised seven early-stage populations enriched for xylem precursors (53.4%) and stele/vasculature initials (11.1%) (**Table S16**). Marker projections reinforced this stage separation (**Fig. 6d**): *LEA3* was broadly enriched in SlAR vascular-associated clusters, whereas SBRL-a key regulator of SBR initiation-remained largely confined to early SBR populations (**Fig. S13; Table S17**). *DOF11* displayed an intermediate distribution with moderate enrichment in SlAR, consistent with its association with the *LEA3*-linked vascular axis.

**Figure 6.**
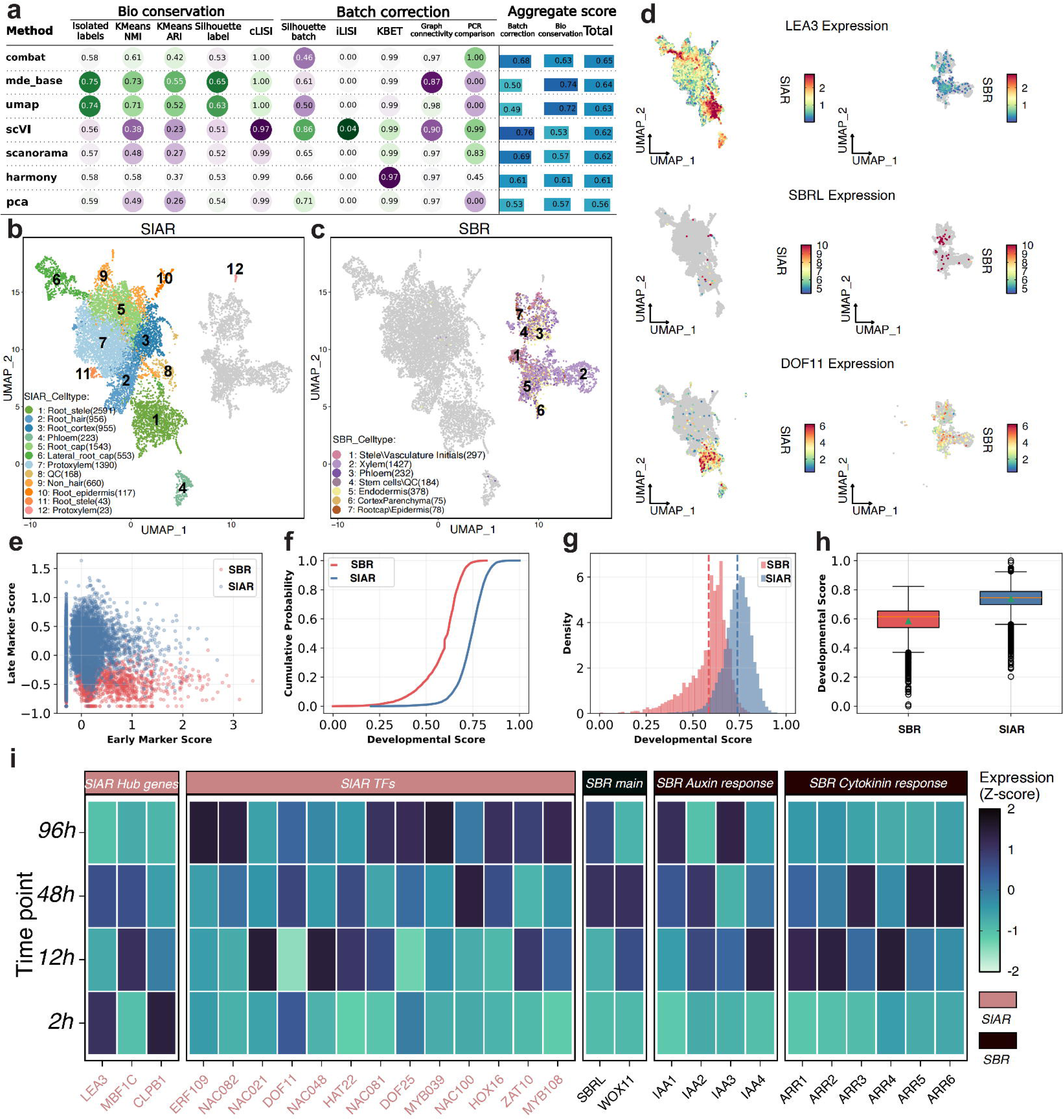
Integration of SlAR and SBR single-cell transcriptomes defines developmental stages of post-embryonic root formation. **(a)** scib-metrics benchmarking of seven batch-correction strategies for integrating SlAR and SBR datasets. **(b,c)** UMAP visualizations of the ComBat-integrated embedding for SlAR and SBR cells, colored by annotated cell types. **(d)** Marker projections across the integrated embedding, highlighting *LEA3*, *DOF11*, and SBRL expression patterns. **(e)** Scatter plot comparing early-stage and late-stage marker scores for individual cells. **(f)** Cumulative distribution functions of the composite developmental score in SBR and SlAR. **(g)** Histogram of developmental score distributions. **(h)** Box plot summarizing score separation between datasets with statistical testing. (i) Bulk RNA-seq heatmap of 28 regulatory genes across four time points (2 h, 12 h, 48 h, 96 h post-excision); expression is Z-score normalized.

To quantify developmental position independent of cluster labels, a composite developmental score was constructed by integrating six early-stage markers (*SBRL*, *PLT*, *WOX5*, *ARF7*, *PIN1*, *IAA1*) with 14 late-stage markers representing differentiated tissues (**Table S19**). Higher scores correspond to more mature states by design. This metric separated the two atlases with minimal overlap: SBR cells occupied lower scores (mean = 0.585 ± 0.108), whereas SlAR cells shifted to higher ranges (mean = 0.738 ± 0.076) (Mann-Whitney U test p < 0.001; Cohen’s d = 1.82; Fig. 6e-h; **Tables S17, S20**).

These results quantitatively place SBR within early primordium specification and SlAR within later tissue differentiation.

Bulk RNA-seq profiling of SlAR samples across four time points (2 h, 12 h, 48 h, 96 h post-excision; three biological replicates each; **Table S21**) provided temporal resolution complementary to the single-cell staging (**Fig. 6i**). Across 28 regulatory genes spanning hdWGCNA hubs, MINI-EX-predicted TFs, and SBR pathway components (**Table S22**), LEA3 uniquely maintained high expression from the earliest sampled time point (2,564 CPM at 2 h) through later stages, with biphasic dynamics peaking at 2 h and 48 h. In contrast, *SBRL* and *WOX11* showed progressive induction from low baseline, while *NAC021*, *HAT22*, and *ERF109* exhibited stage-restricted activation. Together, cross-atlas staging and bulk temporal profiles distinguish *LEA3* from canonical initiation markers and position it as a sustained vascular program associated with AR progression from initiation toward maturation.

### Single-cell atlas integration across eight species positions tomato AR founder vasculature with woody-associated transcriptional programs

The sustained *LEA3* signal identified in vascular AR states prompted a lineage-of-origin question: whether the same marker also delineates the stem vascular cells that give rise to AR founder populations. RNA in situ hybridization on stem sections across four time points supported this continuity. *LEA3* transcripts were detectable in vascular parenchyma cells by 2 h after excision, intensified by 12 h coincident with primordium initiation, and persisted through 48-96 h when visible AR emergence occurred (**Fig. 7a**), indicating that *LEA3* marks stem vascular domains before and during their transition into AR primordia.

**Figure 7.**
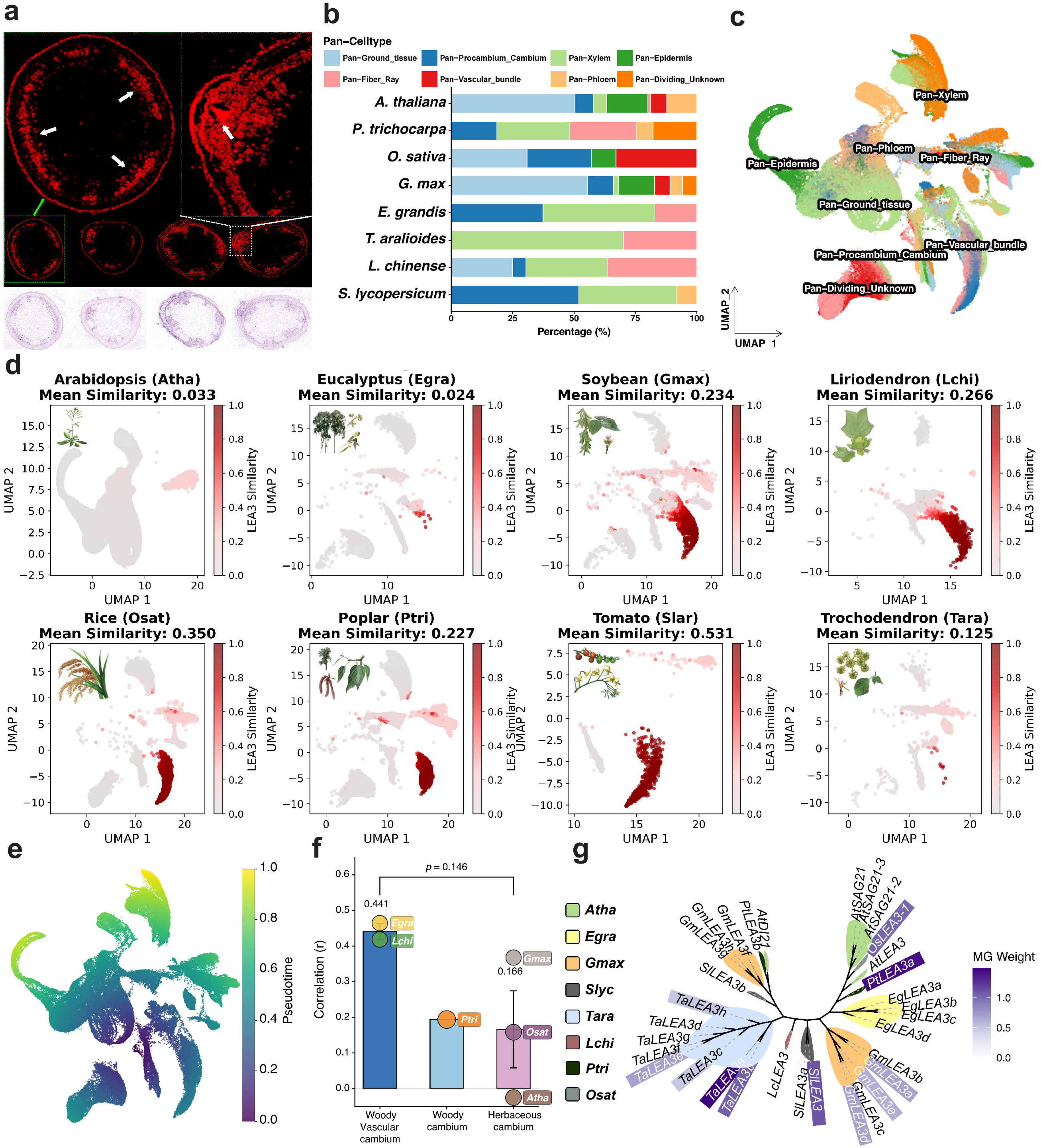
Cross-species stem integration links tomato AR founder vasculature to woody-associated vascular transcriptional programs. **(a)** *LEA3* RNA in situ hybridization on stem sections at 2, 12, 48, and 96 h after cutting. **(b)** Distribution of pan–cell type categories across eight plant species following SATURN integration. **(c)** SATURN-integrated embedding showing eight pan–cell types and the mapping of tomato AR-initiating vascular populations. **(d)** MG_420 (*LEA3*-containing macrogene) expression similarity across species projected onto the integrated space. **(e)** Pseudotime landscape projected onto the integrated embedding. **(f)** Cross-species similarity of tomato Pan-Procambium_Cambium transcriptomes relative to corresponding lineages. **(g)** Phylogenetic tree of *LEA3* homologs annotated with MG_420 weights.

Because AR founder cells are recruited from stem vasculature, the earliest AR-associated populations in the integrated atlas—Pan-Procambium_Cambium, Pan-Xylem, and Pan-Phloem—represent intermediates between stem vascular identity and differentiated root tissues. To place these founder-associated states in a broader phylogenetic context, tomato AR-initiating cells were integrated with stem single-cell transcriptomes from seven additional species (118,730 cells total) using SATURN (**Fig. 7b, c; Table S23**). The resulting embedding resolved eight pan-cell types (**Table S24**) and revealed systematic composition differences across lineages: woody species were enriched for Pan-Fiber_Ray and Pan-Xylem and lacked Pan-Cortex and Pan-Epidermis, whereas tomato founder-associated cells mapped predominantly to three vascular populations—Pan-Procambium_Cambium (51.9%), Pan-Xylem (40.0%), and Pan-Phloem (8.2%) (**Fig. 7c; Fig. S14; Table S25**).

Mapping tomato *LEA3* into SATURN’s protein embedding space identified MG_420 as the *LEA3*-containing macrogene (weight = 1.11), comprising 100 genes across the eight species (**Fig. S15; Table S26**). OrthoFinder-identified *LEA3* orthologs were independently assigned to the same macrogene, supporting cross-species homology at the macrogene level. Notably, expression similarity patterns for MG_420 differed markedly across species: tomato MG_420 correlated with rice (*r* = 0.350) and Liriodendron (*r* = 0.266) but showed near-zero similarity to *Arabidopsis* (*r* = 0.033) (**Fig. 7d; Table S27**). Consistent with this, pseudotime projection placed tomato vascular cells at intermediate differentiation states (mean PT = 0.40), between protoxylem and phloem trajectories (**Fig. 7e; Table S28**). A broader comparison anchored on Pan-Procambium_Cambium further reinforced this asymmetry: tomato founder-associated vascular transcriptomes were most similar to woody dicots, including *Liriodendron* (*r* = 0.46) and *Eucalyptus* (*r* = 0.42), while exhibiting near-zero correlation with Arabidopsis (*r* = −0.02) (**Fig. 7f**).

The *LEA3* regulatory axis showed parallel structure within tomato founder-associated vasculature. *DOF11*—the inferred upstream regulator of *LEA3* in vascular contexts (**Fig. S16**)—displayed cross-species similarity patterns biased toward woody lineages (**Fig. S7a; Table S29**). Within tomato vascular populations, *DOF11* and *LEA3* were strongly coexpressed (*r* = 0.995) with overlapping high-expression domains (**Table S30**). Phylogenetic reconstruction of 32 *LEA3* homologs further contextualized macrogene contribution across species: genes with the highest MG_420 weights—including *Trochodendron TaLEA3b* (1.49), *Populus PtLEA3a* (1.44), and tomato *SlLEA3* (1.11)—were distributed across woody and herbaceous lineages, whereas Arabidopsis *AtLEA3* carried a substantially lower weight (0.44) (**Fig. 7g; Tables S31–S32**). Together, cross-species integration and *LEA3*-*DOF11* centered mapping place tomato AR founder vasculature within a transcriptional neighborhood more closely aligned with woody dicot vascular programs than with *Arabidopsis* (**Fig. S17, S18**).

## Discussion

The developmental transition from differentiated stem tissue to a functional AR represents one of the most striking manifestations of plant cellular plasticity. Unlike embryonic root formation, which proceeds along a well-defined stem cell lineage established during embryogenesis, AR organogenesis requires mature cells to reacquire developmental competence—a process traditionally framed as dedifferentiation followed by reconstitution of a root stem cell niche (Steffens & Rasmussen, 2016; Damodaran & Strader, 2024). Our integrative analyses support a conceptual reframing with broader implications for post-embryonic organogenesis: the boundary between differentiated identity and stemness is not categorical but conditional, with differentiated vascular cells retaining latent stem cell–like transcriptional states that can be rapidly mobilized. By integrating single-cell transcriptomics, cross-dataset developmental staging, and cross-species evolutionary comparisons, we provide evidence that AR competence reflects activation of pre-existing transcriptional potential rather than de novo reconstruction of a stem cell program.

### Vascular tissues as the developmental epicenter: evidence from transcriptional architecture and cross-dataset integration

Our single-cell atlas identifies vascular tissues as the transcriptional epicenter of AR competence. Root stele and phloem cells exhibit the highest CytoTRACE ordering scores, indicating elevated transcriptional diversity associated with high developmental potential (Gulati *et al*., 2020). This observation aligns with emerging concepts in vascular cambium biology, in which the cambium is increasingly viewed as a dynamically organized system comprising stem cells, organizer cells, and transit-amplifying populations rather than a homogeneous meristem (Wybouw *et al*., 2024). Lineage tracing studies demonstrate that a single bifacial stem cell within each radial file generates both xylem and phloem derivatives (Smetana et al., 2019; Shi et al., 2019), and that its position can shift in response to hormonal cues (Mäkilä et al., 2023). Such positional plasticity provides a mechanistic precedent for AR initiation, suggesting that the molecular machinery enabling stem cell repositioning during secondary growth may be redeployed to reacquire stem cell–like states upon wounding.

Consistent with this view, hdWGCNA analysis revealed a modular transcriptional architecture selectively enriched in vascular tissues. Five co-expression modules (4, 6, 7, 8, and 11) were preferentially active in root stele cells, with module 6 centered on *LEA3* as the principal hub gene. This module bridges classical stress-responsive genes (including *CLPB1* and *ERF109*) with developmental regulators, indicating that AR organogenesis exploits regulatory networks that are constitutively available in vascular tissues rather than assembled anew. Given the intrinsic sparsity of single-cell RNA-seq data, robust identification of co-expression modules generally requires aggregation-based strategies such as metacell or pseudobulk representations (Liu & Li, 2025). The stable detection of vascular-specific modules in our analysis is therefore consistent with previous demonstrations that hdWGCNA can recover biologically meaningful co-regulatory relationships from high-dimensional single-cell data, rather than reflecting artifacts of clustering or aggregation (Mallick *et al*., 2022; Morabito *et al*., 2023).

### Cross-dataset developmental staging positions *LEA3* as a constitutive vascular identity marker

Integration with the shoot-borne root (SBR) atlas (Omary *et al*., 2022) enabled precise developmental staging across complementary post-embryonic rooting contexts. The integration of time-lapse imaging with single-cell RNA-seq has revealed distinct temporal phases—from wound response through organogenesis—during de novo root regeneration (Liu *et al*., 2022). Our cross-dataset strategy extends this paradigm to tomato AR formation. Systematic benchmarking identified ComBat as optimal (scib-metrics score = 0.65), addressing the central challenge that naive integration can conflate batch effects with biological differences (Passalacqua & Gillis, 2024). The integrated analysis revealed clear compositional differences: SlAR was dominated by mature root stele (28.0%) and protoxylem (15.0%), whereas SBR was enriched for xylem precursors (53.4%).

A composite developmental score integrating six early-stage markers (*SBRL*, *PLT*, *WOX5*,

*ARF7*, *PIN1*, *IAA1*) and fourteen late-stage markers demonstrated minimal overlap between datasets (Cohen’s d = 1.82). This marker-based staging draws on established paradigms: PLT transcript levels peak at earliest pseudotime points and rapidly decrease during differentiation (Blob *et al*., 2018), while *WOX5* expression is progressively terminated during root-to-shoot conversion (Cui *et al*., 2024). Within this framework, *LEA3* emerged as uniquely informative.

Whereas SBRL showed 5.7-fold enrichment in early SBR cells, *LEA3* displayed inverse enrichment in mature SlAR cells (61.9% versus 12.2%). Bulk RNA-seq confirmed that among 28 key regulators, *LEA3* was the only gene maintaining high expression from 2h through development, with biphasic peaks at 2h and 48h—contrasting sharply with classical stage-restricted markers like *PLT1/2* that decline during differentiation (Maruyama & Ikeuchi, 2025).

This sustained expression pattern positions *LEA3* as a constitutive vascular identity marker rather than a developmental switch. Stage-specific factors like *PLT* and *WOX5* establish transient transcriptional states that are subsequently resolved (Shimotohno *et al*., 2018), whereas constitutive markers may define the permissive cellular context within which these switches operate. *LEA3*’s persistence throughout AR development—from initial wound response through tissue differentiation—suggests it marks a stable vascular identity underlying, rather than driving, developmental progression.

The *DOF11*-*LEA3* regulatory axis: an ancient vascular identity module with stress signal integration

The identification of *DOF11* as the predominant upstream regulator of *LEA3* has implications extending beyond tomato AR development. DOF transcription factors constitute a plant-specific family with remarkable functional diversification: the same DNA-binding specificity has been co-opted for seed development, guard cell function, and vascular patterning (Zou & Sun, 2023; Hunziker & Greb, 2024; Sun *et al*., 2025). Within vascular contexts, the PEAR proteins trigger periclinal divisions by controlling *SMXL3*, a key phloem regulator (Hunziker & Greb, 2024), while *HCA2/DOF5.6* coordinates cambial activity (Guo *et al*., 2009; Zhang *et al*., 2022). This convergence across paralogs suggests deeply conserved vascular functions.

Our network analysis places *DOF11* at the regulatory apex: among all transcription factors evaluated by MINI-EX, *DOF11* achieved the highest global Borda score (570) and strongest interaction weight with *LEA3* in phloem (46.4). This network positioning echoes the broader pattern of DOF proteins functioning as transcriptional hubs through protein-protein interactions. DOF proteins contain bifunctional domains—a conserved N-terminal zinc finger for DNA binding and a non-conserved C-terminal domain enabling interactions with diverse transcription factor families including bZIP, MYB, and WRKY proteins (Manna *et al*., 2021). The *Arabidopsis* DOF protein OBP1 exemplifies this hub function by enhancing bZIP binding to DNA targets (Xu *et al*., 2025), while barley BPBF interacts with GAMYB to activate downstream genes (Su *et al*., 2021). *DOF11*’s apex position in our regulatory network thus reflects the inherent capacity of DOF proteins to integrate multiple transcriptional inputs.

The *LEA3* promoter architecture provides mechanistic insight into signal integration. Four ABREs and one AuxRE (ratio = 4.0) create a regulatory logic enabling ABA-mediated stress signals to converge with auxin-mediated developmental cues. This architecture reflects a broader pattern: composite AuxRE elements frequently incorporate ABRE-like motifs as 5’-flanking or overlapping sequences, with coupling motifs typically positioned within 50 nucleotides to form functional bipartite elements (Berendzen *et al*., 2012). Such bipartite promoters enable transcription factors with dual DNA-binding specificity—like ABI3-like factors that recognize both ABA and auxin response elements—to mediate hormone crosstalk

(Suzuki *et al*., 2001; Liu *et al*., 2013). Indeed, DOF proteins themselves participate in ABA-GA crosstalk: *Arabidopsis DAG1* interacts with the DELLA protein GAI to modulate GA biosynthesis (Boccaccini *et al*., 2014), while the *RGL2-DOF6* complex regulates seed dormancy through GATA transcription factor activation (Ravindran *et al*., 2017; Gao *et al*., 2025). At 48h—coinciding with primordium emergence—*LEA3* expression was strongest under IAA+ABA co-treatment, with synergistic effects exceeding either hormone alone. This positions LEA3 as a molecular node where the *DOF11*-initiated vascular identity program integrates with environmental stress monitoring, consistent with the emerging view that auxin-ABA crosstalk is fundamental to growth-defense trade-offs during organogenesis (Müller & Munné-Bosch, 2021).

Cross-species integration reveals evolutionary alignment with woody vascular programs The challenge of cross-species transcriptomic comparison lies not merely in technical integration but in the fundamental question of homology. Traditional approaches rely on one-to-one orthologs, yet frequent whole-genome duplications in plants create complex many-to-many relationships, and genes with strict one-to-one correspondence are enriched for essential genes under single-copy control rather than representing the transcriptome broadly (Church *et al*., 2024). SATURN addresses this by leveraging protein language models (ESM2) to embed genes in shared functional space, grouping them into “macrogenes” based on protein embedding similarity (Rosen *et al*., 2024). This approach parallels recent advances demonstrating that protein structure-based comparisons can reveal functional conservation obscured by sequence divergence—a principle validated across diverse taxa from *Arabidopsis* to maize despite 160 million years of evolutionary separation (Zhai *et al*., 2025).

Our cross-species integration of 118,730 vascular cells from eight species revealed an unexpected pattern: tomato AR-initiating cells showed preferential transcriptomic similarity to woody dicots (*Liriodendron*, *r* = 0.46; *Eucalyptus*, *r* = 0.42) rather than the herbaceous model *Arabidopsis* (*r* =-0.02). This finding resonates with emerging evidence that xylem developmental programs are not uniformly conserved. Cross-species single-cell analysis of xylem development has revealed that while ray lineages are highly conserved across angiosperms, fusiform lineages—comprising vessel elements and fibers—exhibit substantial variability between core eudicots, basal eudicots, and magnoliids (Tung *et al*., 2023). The tomato-woody similarity we observed may reflect shared fusiform developmental programs that diverged in *Arabidopsis*.

Why might *Arabidopsis* represent a derived rather than ancestral state for AR competence? Multiple lines of evidence suggest regulatory rewiring. Transcription factors functioning as positive AR regulators in woody species act as negative regulators in *Arabidopsis* (Kidwai et al., 2023), and AR formation capacity is notably constrained in *Arabidopsis* compared to species dependent on vegetative propagation (Druege *et al*., 2019). The vascular cambium itself differs fundamentally: trees maintain seasonal oscillations from dozens of cambial cell layers to few, requiring elaborate stem cell maintenance programs (Wybouw *et al*., 2024) that *Arabidopsis* minimal secondary growth does not demand. Tomato, while herbaceous, undergoes substantial secondary growth and may retain ancestral cambial regulation—a pattern consistent with the observation that “herbaceous” and “woody” classifications obscure vast anatomical variation in secondary growth capacity (Spicer & Groover, 2010).

The phylogenetic distribution of macrogene weights illuminates the evolutionary trajectory of vascular-to-root developmental programs. Our comparison spans species representing over 100 million years of angiosperm evolution: *Liriodendron* (magnoliid, diverged ∼170–145 Ma) retains ancestral vascular organization predating the monocot-eudicot split (Chen *et al*., 2019;

Dong *et al*., 2022), while *Trochodendron*—despite its vessel-free wood, an evolutionarily reversed character—maintains tracheids transcriptomically similar to vessel elements, suggesting conservation of ancestral water transport programs (Tung *et al*., 2023). Critically, secondary growth from bifacial vascular cambium represents the ancestral condition in seed plants, with herbaceous habits constituting derived reductions (Spicer & Groover, 2010; Cunha Neto, 2023). This evolutionary framework reframes our macrogene results: the high MG_420 weights in basal angiosperms (*Trochodendron*, 1.49; *Liriodendron* implicit in cross-species correlation) and woody dicots (*Populus*, 1.44) likely reflect retention of ancestral vascular identity programs, whereas the markedly reduced weight in *Arabidopsis* (0.44) represents derived loss accompanying its transition to a rapid annual lifecycle with minimal cambial activity. Tomato, though classified as herbaceous, undergoes substantial secondary growth and accordingly retains ancestral-like *LEA3* expression (weight = 1.11). This pattern suggests that the capacity for vascular-derived adventitious rooting is not a specialized adaptation but an ancestral competence embedded in cambial stem cell programs. The reduced *LEA3* weight in *Arabidopsis* may reflect a derived trade-off: recent evidence demonstrates that regeneration capacity declines with plant maturation through salicylic acid-mediated defense enhancement, revealing a fundamental tension between regenerative flexibility and rapid lifecycle completion (Xu & Yang, 2025). Annual species like *Arabidopsis*, which allocate resources predominantly toward rapid reproduction rather than sustained vegetative competence (Friedman, 2020; Zhao *et al*., 2023), may have selectively reduced these ancestral regenerative programs.

## Acknowledgements

This work was supported by National Natural Science Foundation of China (32560738), Research and Utilization Center for Qinghai-Tibet Plateau Germplasm Resources (2025 Program).

## Author contributions

Conceptualization, SY and RY; Formal analysis, SY and XJ; Funding acquisition, SY and QZ; Investigation, CS and XS; Methodology, SY, SC and CS; Writing—original draft, SY, XJ; Writing—review & editing, SY, RY and CS. All authors have read and agreed to the published version of the manuscript.

## Data availability

The tomato AR single-cell RNA sequencing data generated in this study have been deposited in databases. The raw sequencing data are available in the NCBI Sequence Read Archive (SRA) under BioProject (https://www.ncbi.nlm.nih.gov/sra/PRJNA1218210). The same dataset has also been deposited in the National Genomics Data Center (NGDC) under Project (PRJCA035692). The processed single-cell data, including Seurat objects, analysis scripts, and associated files are available through Figshare (https://figshare.com/projects/Single-cell_Resolution_Analysis_Reveals_Tomato_adventitious_root_Development_and_Evolution/2 36603). All analysis code and supplementary data used in this study are also available in the same Figshare repository.

## Supporting Information

Additional Supporting Information may be found online in the Supporting Information section at the end of the article.

**Method. S1** Optimized Protoplast Isolation Protocol for Single-cell RNA Sequencing of Tomato Ars.

**Method. S2 Promoter Cis-Regulatory Element Analysis**

**Method. S3 Quantitative Real-Time PCR (qPCR) Validation**

**Method. S4** Cross-Dataset Integration and Batch Correction

**Method. S5** Bulk RNA-Seq Analysis

**Method. S6** SATURN Cross-Species Single-Cell Integration

**Fig. S1** Comparison of different resolution parameters for the RNA_snn_res method in Seurat.

**Fig. S2** Performance comparison of Random Forest and SVM models in cell type classification.

**Fig. S3** Detailed comparison of SVM and Random Forest classification performance across root cell types.

**Fig. S4** Heatmap of tomato adventitious root marker genes identified by SPmarker.

**Fig. S5** Distribution of 10 topics across different cell types.

**Fig. S6** Network visualization of topic relationships and their associated GO terms.

**Fig. S7** Determination of optimal soft power threshold for hdWGCNA network construction.

**Fig. S8** Spatial distribution maps of module expression based on uCell scores.

**Fig. S9** Cis-regulatory elements identified in the 2-kb promoter regions of *LEA3* (a), *CLPB1* (b), and *MBF1C* (c).

**Fig. S10** Phylogenetic analysis of *LEA* family proteins across plant species.

**Fig. S11** Phylogenetic analysis of *CLPB* family proteins across plant species.

**Fig. S12** Cell type-specific transcription factor enrichment and expression analysis.

**Fig. S13** Differential gene expression patterns between mature SlAR maintenance and early SBR initiation stages.

**Fig. S14** Distribution of tomato vascular cell types on SATURN umap

**Fig. S15** UMAP overview of pan-cell type annotation and expression patterns of *DOF11* and *LEA3* in the cross-species single-cell dataset.

**Fig. S16** MG_382 (*DOF11*-containing macrogene) expression similarity across species projected onto the integrated space.

**Fig. S17** Pairwise cell composition similarity matrix across eight plant species.

**Fig. S18** Phylogenetic analysis of DOF11 transcription factor family with macrogene weights across eight plant species.

**Table. S1** Mapping statistics for tomato AR single-cell sequencing data.

**Table. S2** Sequencing data quality metrics from Cell Ranger pipeline.

**Table. S3** Cell and gene counts per cluster after quality control filtering.

**Table. S4** Marker genes for each cell type with differential expression statistics.

**Table. S5** Target genes and module membership from hdWGCNA analysis.

**Table. S6** Cis-regulatory elements in 2 kb promoter regions of hub genes.

**Table. S7** Transcription factor binding motif enrichment summary.

**Table. S8** qPCR validation data across hormone treatments and time points.

**Table. S9** MINI-EX output of transcription factor–target gene regulatory pairs.

**Table. S10** Gene ID to gene name conversion.

**Table. S11** Network topology statistics for TF-TG relationships.

**Table. S12** Regulatory correspondence between MINI-EX TFs and hdWGCNA hub genes.

**Table. S13** Integration parameters for SlAR and SBR datasets.

**Table. S14** scib-metrics benchmark of batch correction methods.

**Table. S15** Cell type composition of the SlAR dataset.

**Table. S16** Cell type composition of the SBR dataset.

**Table. S17** Developmental score statistics comparing SlAR and SBR.

**Table. S18** Marker gene expression comparison between datasets.

**Table. S19** Early-stage and late-stage marker genes for developmental scoring.

**Table. S20** uCell module enrichment scores across cell types.

**Table. S21** Bulk RNA-seq expression of 28 key genes across four time points.

**Table. S22** CPM values and temporal dynamics of key regulatory genes.

**Table. S23** Cell counts and quality metrics for eight-species SATURN integration.

**Table. S24** Pan-celltype definitions from cross-species integration.

**Table. S25** Species-specific pan-celltype distribution.

**Table. S26** *LEA3* orthologs within macrogene MG_420 across eight species.

**Table. S27** *LEA3* cross-species expression correlation coefficients.

**Table. S28** Pseudotime values for cross-species trajectory analysis.

**Table. S29** *DOF11* cross-species expression correlation coefficients.

**Table. S30** *DOF11*-*LEA3* coexpression statistics in tomato vascular cells.

**Table. S31** Complete gene membership of macrogene MG_420.

**Table. S32** *DOF11* orthologs within macrogene MG_382 across eight species.

